# Phenotypic reversion and target prioritization for cellular inflammation via representation learning with foundation models

**DOI:** 10.64898/2026.03.04.709607

**Authors:** Daniel R. Wong, Mary Piper, Jiao Qiao, Max Russo, Pierre Jean, Djork-Arné Clevert, Jason D. Arroyo, Evanthia Pashos

## Abstract

The identification of genetic perturbations that can reverse disease-associated cellular phenotypes toward a healthy state is a central challenge in early drug discovery. We present a proof-of-concept framework leveraging single-cell foundation models (scFMs) and a large-scale Perturb-seq dataset to prioritize targets for phenotypic reversion of cellular inflammation. We incorporated both basal and proinflammatory signaling conditions, specifically stimulation with interleukin-1 beta (IL-1β) and tumor necrosis factor alpha (TNF-α), to assess whether atherosclerotic disease-relevant stimulation improved identification of genes and pathways critical to disease progression. Our dataset comprised 864,115 endothelial cells subjected to 1,740 unique genetic perturbations. Through having both conditions, we identified targets that exhibited differential effects on gene expression dependent on cellular state. Using scFMs, we embedded single-cell transcriptomes into high-dimensional latent spaces and ranked perturbations by their ability to shift the inflammatory transcriptomic profile toward that of untreated controls. Benchmarking against both annotated gene sets and expert-curated targets, we found robust enrichment for known regulators of inflammation and biologically relevant targets, despite these models having no prior information about these targets. Importantly, including both basal and proinflammatory conditions improved identification of inflammation-associated targets compared to using just the basal condition. This underscored the value of incorporating disease-relevant stimulations in perturbation experiments. Our results highlight the utility of scFMs for data-driven target nomination, emphasize the role of cellular state in regulatory responses, and provide a scalable, model-agnostic approach for ML-guided target discovery. This work offers a valuable community resource for advancing biological understanding of inflammatory-associated disease.

## Introduction

Transcriptomics^1–3^ is a fundamental area of biology that provides a window into the biological state of tissues. In recent years, single-cell sequencing has revolutionized the field by providing a granular snapshot of the transcriptome on a per cell basis^4,5^. Combining this technology with perturbation via small molecules or genetic alterations like CRISPR^6^ provides an unprecedented opportunity to study disease biology and how different perturbagens can potentially provide remediation. Consequently, technologies like Perturb-seq^7^, which measures the post-perturbation transcriptome of single cells, provide a unique opportunity in the drug discovery space via both its scalability and granularity to measure subtle phenotypic changes.

One interesting area of inquiry concerns the idea of “phenotype reversion,”^8^ which explores whether certain perturbations can revert a diseased state phenotype to one resembling a healthy state. For early discovery, such a readout can be a multitude of phenotypes, such as high-resolution images^9–12^, proteomics^13^, or single-cell RNA-sequencing data (scRNA-seq). There has been some success in these areas for early discovery^8,14–17^. With the recent rise of Machine Learning (ML) capabilities, and in particular foundation models^18,19^, there is an opportunity to combine the technologies of scRNA-seq and ML with rapidly increasing interest and innovation in this area^20–25^. Through large-pretraining on diverse datasets, scFMs have shown promise in creating meaningful representations of cells which can be used for various downstream tasks of interests^26^. Although tasks such as cell type annotation, prediction of the post-perturbation transcriptome^20,23,27,28^, and forecasting differential gene expression^25,29,30^ have all undergone much inquiry, exploring the propensity of scFMs to assist with target nomination has remained a relatively unexplored area. Associating the knockdown of specific targets to desirable phenotypes is an initial yet essential task in evaluating a target’s potential for therapeutic intervention.

We generated a Perturb-seq dataset with the goal of identifying those genes whose inhibition could reverse proinflammatory signaling and potentially inhibit inflammation-driven atherosclerotic plaque initiation or progression. Proinflammatory signaling cascades associated with increased levels of IL-1β and TNF-α mimic the inflammatory environment present within the plaque regions of the arterial wall in atherosclerotic disease^31^. In atherosclerotic plaques, IL-1β and TNF-α signaling results in NF-kB induction and activation of endothelial cells, resulting in increased expression of proinflammatory cytokines and chemokines and up-regulation of adhesion molecule expression, such as ICAM-1. Increased expression of adhesion molecules on the surface of endothelial cells enables monocyte tethering and migration into the vessel wall, which triggers macrophage activation, lipid accumulation, and foam cell formation. This cascade leads to increased release of cytokines, chemokines and reactive oxygen species, driving chronic inflammation through additional monocyte recruitment and plaque progression ^32,33^. Reversing the inflammatory signaling loop in endothelial cells could potentially disrupt atherosclerotic initiation or progression^34,35^.

Here we present a case study and proof-of-concept for phenotype reversion using scFMs in a real-world pharmaceutical target discovery program. With this publication, we release a rich and high quality Perturb-seq dataset for studying inflammation in endothelial cells, benchmark different foundation models on the task of target nomination for phenotype reversion, investigate the benefit of having a specific disease-associated stimulation, and explore specific biological trends in the inflammation response. With the uptake in both excitement and investment in the prospect of foundation models, investigating how these models can affect target nomination is an unchartered but pressing question that must be answered to assess a new avenue for ML in the early discovery phase. We demonstrate how foundation models can create cellular latent representations that can then be used in a model-agnostic framework for target triaging, with validation via enrichment of known targets and pathways.

## Results

### Large Perturb-seq dataset models cellular inflammation

A high-level overview of the study is illustrated in Figure 1. Briefly, we constructed a Perturb-seq dataset to model cellular inflammation, took multiple approaches for ranking targets based on their propensity to revert inflamed cells to a state most similar to basal control, and evaluated the different rankings via enrichment analyses of *a priori* chosen targets and pathways.

**Figure 1:**
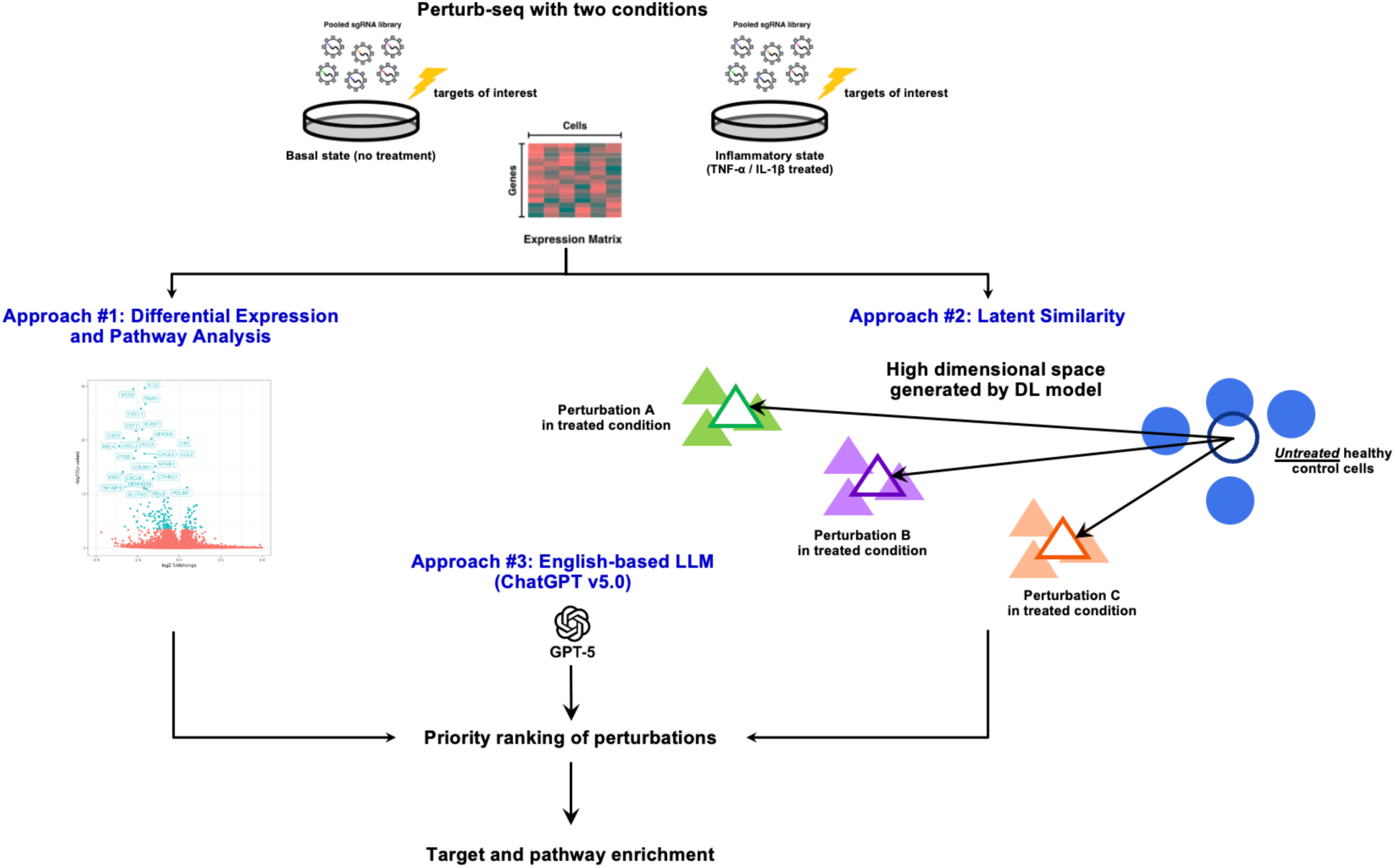
Overview of phenotypic reversion study.

To elucidate inflammatory signaling processes in endothelial cells, we generated a large Perturb-seq dataset using hTERT-immortalized human arterial endothelial cells (TeloHAEC) grown under both basal and inflammatory conditions. We are releasing this unique dataset to encourage the ongoing advancement of ML in early drug discovery and contribute to the growing wealth of publicly available data that can be used for model development, inference, and biological inquiry. The dataset comprised 864,115 teloHAEC cells with expression of 38,606 genes (including both coding and non-coding genes), and 1,740 unique genetic perturbations (of which 870 were pursued due to association with atherosclerotic disease). The perturbed cells were either treated with IL-1β and TNF-α stimulation (which we refer to as the “Treated” or “Inflammatory” condition) or grown in the absence of cytokine stimulation (which we refer to as the “Untreated” or “Basal” condition). From a Uniform Manifold Approximation and Projection (UMAP) visualization of control cell expression for each condition, the inflammatory cytokines exerted a strong effect on transcriptomic expression, with an easily differentiable phenotype (Figure 2A). Moreover, projecting canonical inflammation-relevant genes via UMAP showed transcriptional signs of a differential inflammatory phenotype between the two conditions (Figure 2B). 59% of gene targets exhibited high differential gene expression in one treatment condition but not the other (Figure 2C), revealing how cytokine treatment can largely influence how different targets induce variable transcriptomic changes. Example volcano plots for such targets are shown in Supplemental Figure 1.

**Figure 2:**
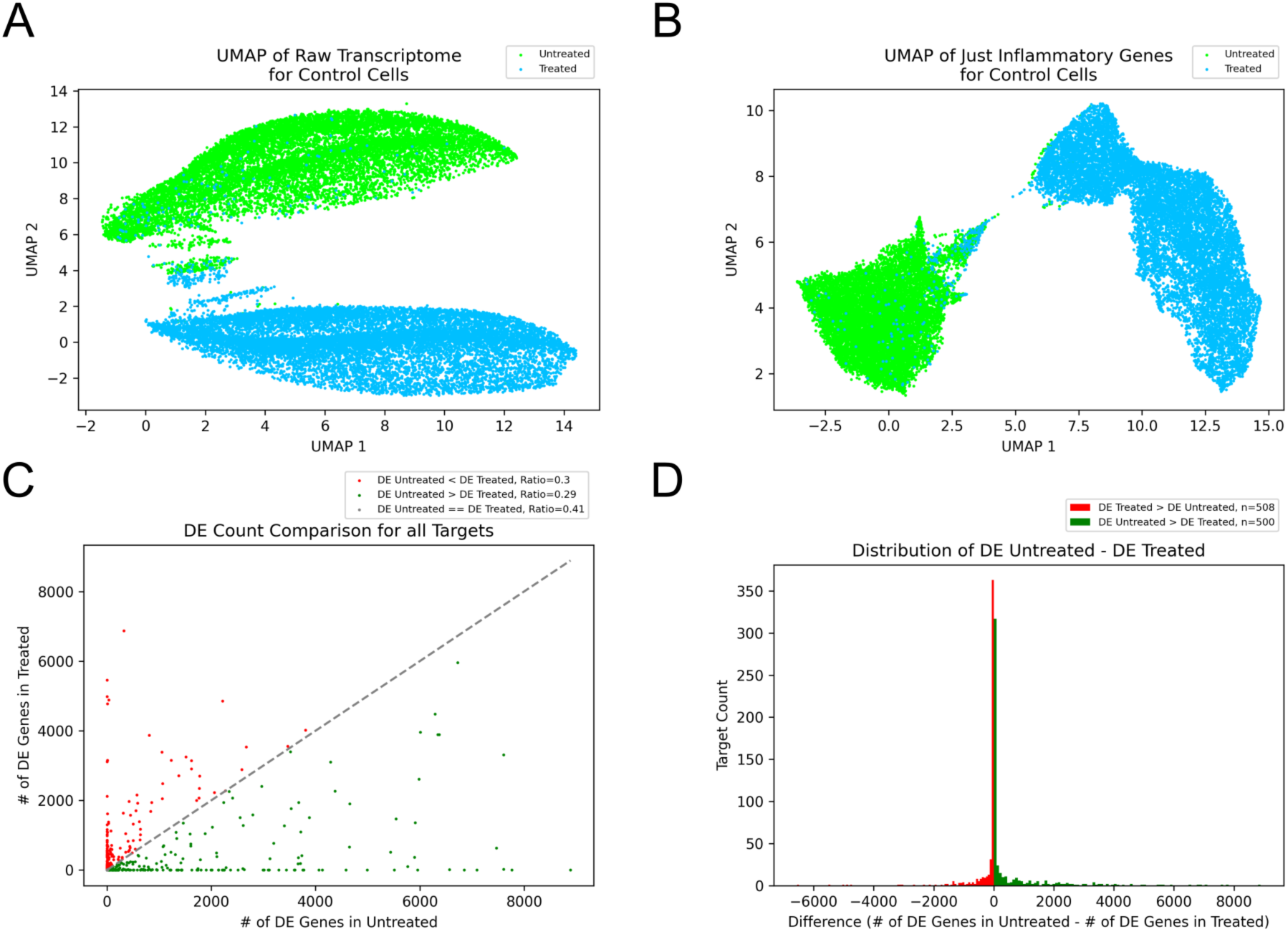
Dataset analysis. (a) UMAP of all control cells (safe-targeting and no-targeting) in both treated (TNF-α and IL-1β) and untreated conditions. (b) UMAP of all control cells over just inflammatory genes. We used the set of genes belonging to the following relevant canonical pathways from MisgDB C2 to compute the UMAP: BioCarta: IL1R Pathway, BioCarta: TNFR1 Pathway, KEGG: Cytokine-Cytokine Receptor Interaction, Reactome: Interleukin 1 Family Signaling, Reactome: Interleukin 1 Signaling, Reactome: TNF Signaling. (c) Target DE in the two conditions. Each point is a gene target. X-axis is the number of DE genes in the untreated condition. Y-axis is the number of DE genes in the treated condition. (d) Histogram of DE Untreated – DE Treated. Each sample is a unique target. Plot excludes targets that had equal number of DE genes in Treated and Untreated.

The dataset had high knockdown efficiency and depth of coverage. The distribution of target gene replicates was similar for each condition, with a median of 188 cells per target for the untreated condition and 194 cells per target for the treated condition. Like most scRNA-seq data, expression of many genes was sparse with the proportion of zero-valued counts in the cell by gene matrix of 87%. Knockdown (KD) efficiency was high, with a KD mean of 72% and a median of 80% of cells (Supplemental Figure 2). There was also a high depth of sequencing per cell, with a median UMI per cell of 16,373.

### Latent similarity approach enriches for positive-control targets

Our goal was to determine which genetic perturbations induced a change from the inflammatory state induced by TNF-α and IL-1β to one resembling the transcriptome of the untreated basal control, thereby effectively reversing the inflammatory phenotype. We took three different broad approaches to triage our targets: ranking based on differential expression (DE) combined with gene set enrichment analysis, representation learning using scFMs, and English-based LLM application.

For the first approach, we first performed DE analysis between each knockdown target relative to the control guide cells using the Wilcoxon rank sum test. We then performed Gene set enrichment analysis (GSEA) on the signed log-transformed p-values from each DE comparison (Methods). We created two different rankings using this approach: (1) DE (Basal) to model a DE analysis if we only had the basal condition and (2) DE (Inflammatory) to model a DE analysis utilizing information from both conditions. The DE (Basal) ranking was calculated based on the significance of down-regulation of the signaling pathways directly downstream of TNF-α and IL-1β (Methods). For DE (Inflammatory), targets were prioritized if enriched pathways were significant in inflammatory, but not basal conditions, indicating greater specificity for the inflammatory response. DE (Inflammatory)’s top 25 targets and their corresponding normalized enrichment scores for the different pathways are found in Supplemental Figure 3. This first approach, which we denote as the “DE approach,” is a classical analysis that does not rely upon any ML-based workflows.

For the second approach, we created representations for each of our genetic perturbations and the safe-targeting control in the treated condition and a representation for our safe-targeting control in the untreated condition using different scFMs. For comparison, we also used non-ML based representations such as UMAP and the raw integer counts of each transcript as features (Methods). With the motivation of identifying targets that revert an inflammatory state to a basal state, we computed which treated target representations were closest to the untreated safe-targeting control in latent space. We ordinally ranked our perturbations by cosine similarity to the control state, measuring which treated perturbations induced a cellular transcriptomic state most similar to the average of the untreated safe-targeting control. The distribution of cosine similarities was variable between the different methods (Supplemental Figure 4). This second approach is designated as the “latent similarity” approach. As a preliminary test of representation specificity, we determined how well the different scFMs retained true biological signal. We assessed the scFMs’ latent spaces by labeling each cell with their gene target, computing clustering scores based on these labels, and comparing with random and baseline representations. For each treatment condition, all representations (besides the random representation) had a ratio greater than 1.0 between Calinski-Harabasz scores of true vs random labels, indicating some degree of intra-target similarity in their latent space (Supplemental Figure 5).

As a third and final approach, we generated a priority ranking independent of the expression data by prompting ChatGPT for a target ranking given the biological context of the experiment and the goal of phenotype reversion (Methods). This approach did not have access to any numerical data in the experiment and relied solely on insights gleaned during its large pre-training independent of this study.

#### Ranking comparisons

Despite the widely varying approaches to ranking targets, there was fair concordance of target rankings between the DE (Inflammatory) approach and the latent similarity approaches, especially those generated with scFMs. The DE (Inflammatory) approach had a rank-biased overlap (RBO) of 64% for STATE, 52% for scGPT, and 48% for SCimilarity. In the top ten prioritized targets, 50% of the DE (Inflammatory) ranking were shared by STATE as well as scGPT (Figure 3A). Many targets were ranked highly by all the approaches (Figure 3B). The DE (Inflammatory) approach had fair concordance with the scFM rankings, while the DE (Basal) approach was most divergent, providing only 0-1% RBO with any of the other methods, including the DE (Inflammatory) method. Latent similarity via UMAP and ChatGPT also did not yield similar rankings to the scFM and DE (Inflammatory) methods. Although ChatGPT had some common highly ranked targets, the rankings were dissimilar to the other two approaches, which both made their inferences directly from the expression data (Figure 3A-B).

**Figure 3:**
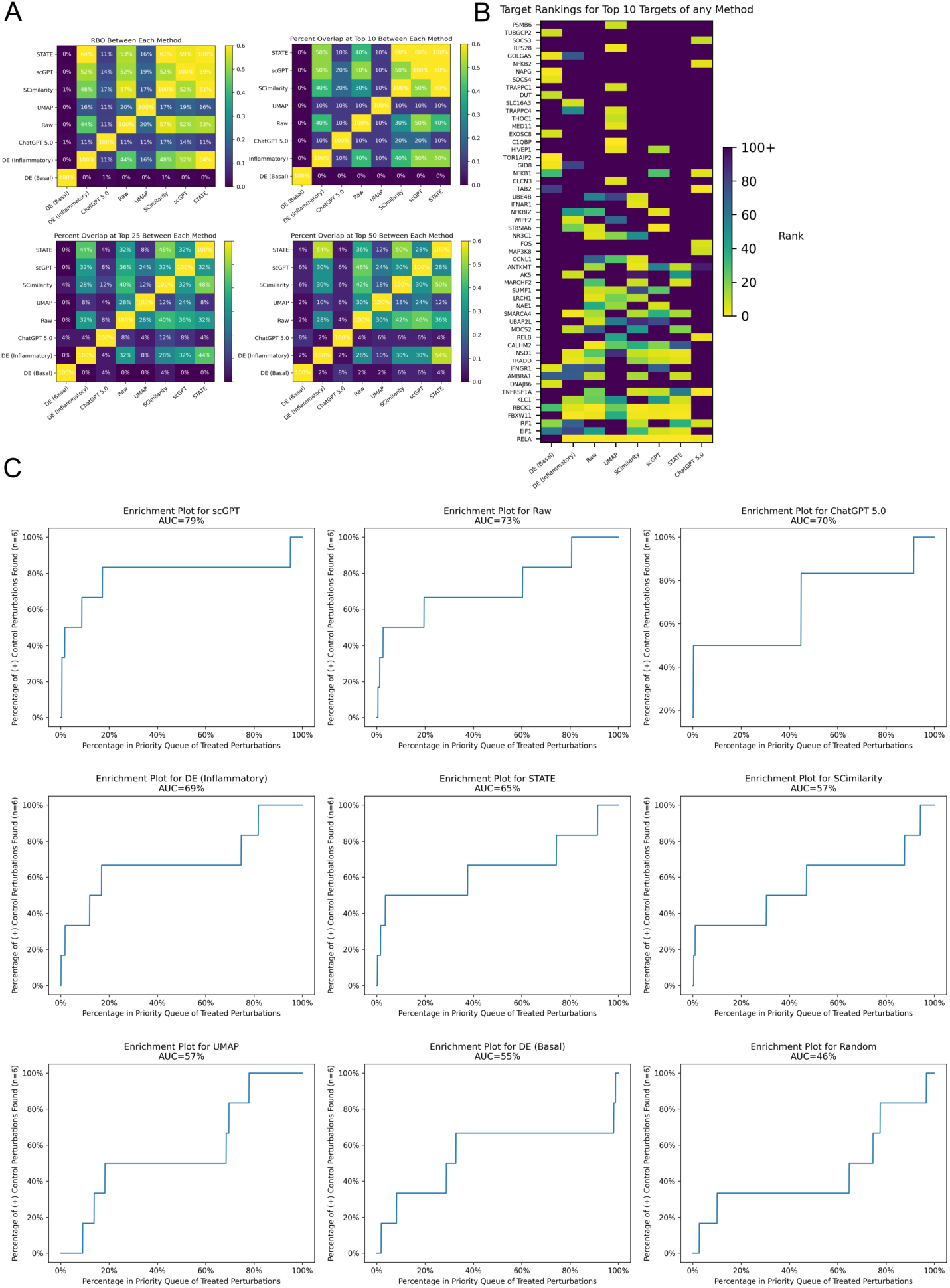
Evaluation of different target rankings. (a) Similarity between the target rankings for different methods. Pairwise rank-biased overlap (RBO) with p=0.90 between each method’s ranked lists. Higher RBOs indicate more similarity, with an exact match having RBO = 1.0. Also shown are percent overlaps at top n for different values of n. (b) Targets part of the top 10 in of any method’s rankings. A target with lower rank is higher priority. (c) Individual enrichment plots for all the methods using the positive-control gene set. AUC = area under the enrichment curve. X-axis is the index in the ranked priority queue expressed as a percent, Y-axis is the percent of positive control targets found at index = x or lower.

#### Positive control enrichment

Once target rankings for the different approaches were established, we asked how biologically sensible they were for potential therapeutic remediation. By the nature of early target discovery, “ground truth” targets that can remediate a disease-associated phenotype are often completely unknown. Hence, to assess a target ranking’s relevance for phenotype reversion, we compared the rankings of different approaches with a set of target genes known to be implicated in the TNF-α and IL-1β signaling cascade. Specifically, we used a small set of genes with strong literature support for which we *a priori* hypothesized their knockdown would suppress the inflammatory phenotype: the major TNF-α receptor (TNFRSF1A), the major TNF-α signaling adaptor (TRADD), and transcription factors known to mediate inflammatory gene expression (JUNB, JUND, NFKB1, NFKB2).

#### These genes served as a positive-control set of targets

We assessed how these different targets were ranked via calculating area under the curve (AUC) for enrichment. The latent similarity approach with scGPT had the highest performance among all approaches (Figure 3C). scGPT target rankings yielded an AUC = 0.79, with positive-control targets ranked as higher priority in scGPT’s rankings compared to any other method. The next highest scoring approach was the latent similarity approach using the raw integer counts as feature embeddings (enrichment AUC = 0.73), followed closely by ChatGPT (enrichment AUC = 0.70) and the DE (Inflammatory) approach (enrichment AUC = 0.69).

We also compared our rankings using the same relevant annotated pathways used to create our DE-approach rankings (Methods). Although the gene sets of these pathways included genes that are broadly related to the TNF-α and IL-1β signaling cascade, they do not reflect the specific context of inflammation reduction. Hence, they may not be the most accurate or specific sets to score our rankings, which were ordered by potential for phenotype reversion. However, these gene sets can illuminate how well the different rankings enrich for known targets that have broad relevance to the TNF-α and IL-1β signaling cascade. The DE-approach rankings yielded the highest AUC for triaging related targets for 5/6 of the biological pathways (Supplemental Figure 6). The targets of the TNFR1 pathway from BioCarta were highly enriched for most of the latent similarity methods (scGPT AUC = 0.91, RAW AUC = 0.85, SCimilarity = 0.82).

### Top targets reveal enrichment of relevant pathways

Once we determined how well the different rankings enriched for targets of interest, we asked how well the top targets of each ranking collectively enriched for biological pathways of interest. This was different than the enrichment of differentially expressed genes typically performed for GSEA analysis (like the one used to construct the DE-approach rankings) and instead focused on possible enrichment of a triaged *target set* for pathways of interest. To determine if the top triaged targets were significantly associated with such pathways, we performed pathway enrichment via Enrichr^36,37^. For each ranking we assessed whether the set of top n gene targets yielded significant enrichment for the different relevant pathways chosen *a priori* (the same ones used to generate the DE-approach rankings). We varied n from top 10 to top 100, and calculated which fraction of the pathways were significantly enriched. Like the target enrichment analysis, scGPT yielded the highest AUC = 0.95 for relevant pathway enrichment. The fraction of recovered relevant pathways varied depending on the rank index, but holistically latent similarity with scGPT resulted in the highest number of relevant pathways, with 100% of relevant pathways recovered from n=30 to n=100 (Figure 4A). Also like the target enrichment analysis (Figure 3), Random embeddings and the under-represented UMAP representation (with just two dimensions) yielded the worst performance (Figure 4A).

**Figure 4:**
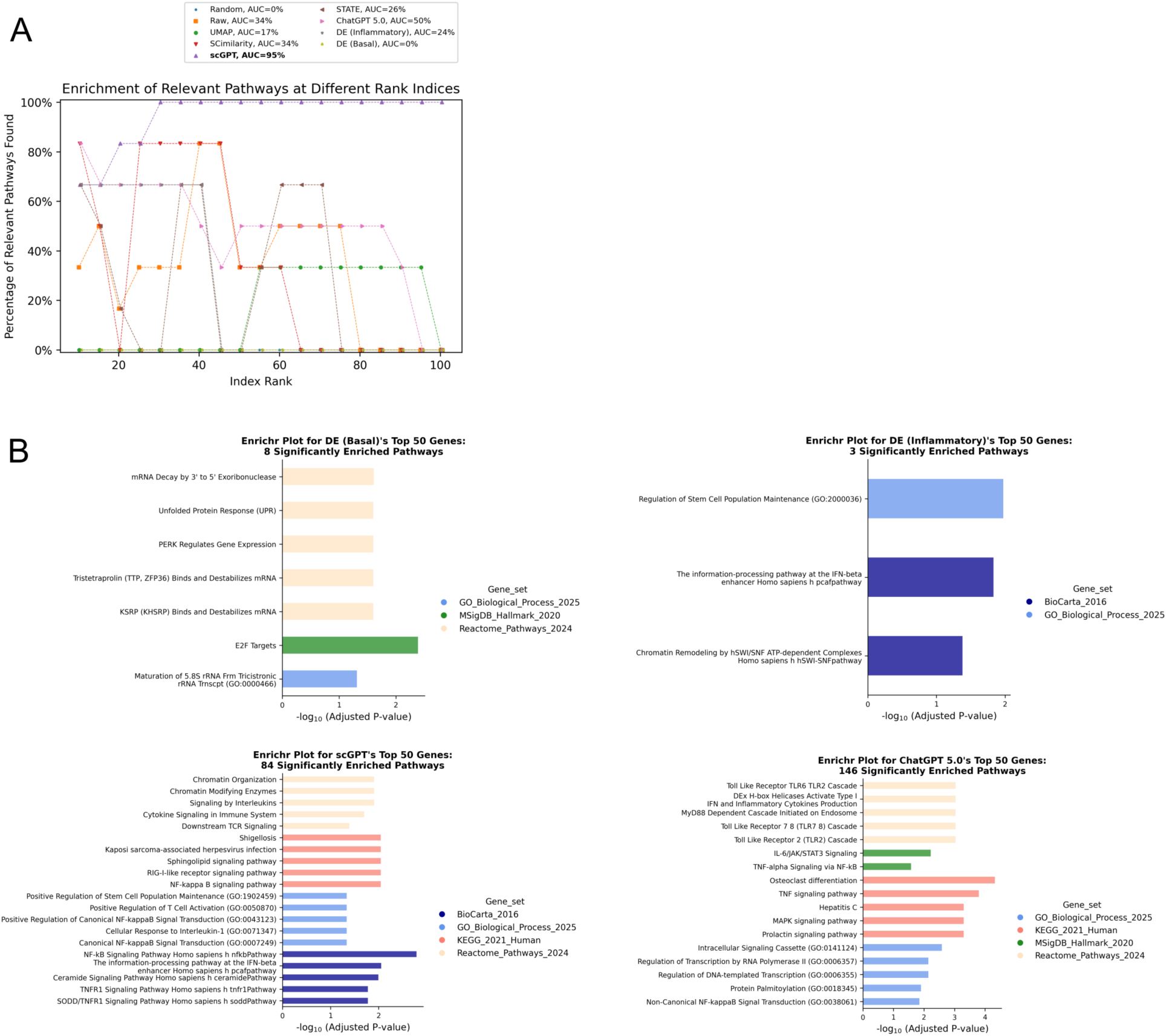
Pathway enrichment for top n ranked targets. (a) Comparison between methods and their propensity to enrich for relevant pathways (same pathways used to construct the DE-approach rankings, see Methods). X-axis is the index in the priority queue, such that targets with rank = 1 to rank = x were included in the input to Enrichr. Y-axis is recall using the relevant pathways as true positives. AUC is the area under the curve for each plot. (b) Significantly enriched pathways using the top 50 targets for DE (Basal), DE (Inflammatory), scGPT, and ChatGPT. We queried five different reference libraries: KEGG_2021_Human, BioCarta_2016,Reactome_Pathways_2024, MSigDB_Hallmark_2020, and GO_Biological_Process_2025. The top five (at most) significant pathways (with p-adjusted < 0.05) of each set are displayed. The total number of pathways achieving significance are reported in the title of each plot.

Instead of focusing on a specific set of relevant pathways, we also performed an enrichment analysis with a broad array of pathways, and tested significance for all pathways in five canonical library annotation sets: KEGG, BioCarta, Reactome, MSigDB, and GO. The DE approach resulted in few significantly enriched pathways, all of which did not appear related to the biological context being studied. In contrast, pathways for the top performing model (scGPT) were sensible and directly related to TNF-α and IL-1β mediated biology despite having no information on pathway to inform target rankings (Figure 4B).

## Discussion

With a growing interest in the development and adoption of ML in the biopharmaceutical space, there is an increasing need not just for new models, but for large and diverse datasets which are of equal importance. From a benchmarking perspective, this dataset that we are releasing has not been used in any training of current foundation models in the field and therefore can be of high utility for benchmarking and testing zero-shot generalization to an unseen context. From a training perspective, new Perturb-seq data is of high value, and can be used for training new models to predict the effects of held-out perturbations and context generalization across diverse cellular states, all of which are experiencing rapid development via collaborative efforts focused on the development of the “virtual cell”^38^. Most ML models require training on large and diverse data to perform well, and we hope that adding our dataset of 864,115 cells with high sequencing depth to the wealth of open-source datasets will facilitate accelerated progress in the field. Having both an inflammatory cytokine condition and genetic perturbations matched in different conditions is less common in Perturb-seq datasets and adds greater biological diversity to the wealth of RNA-seq data. This unique setup provides ample opportunity for learning in a less explored biological space, both through ML and more traditional inquiry. The extensive knockdown of gene targets combined with the high sequencing depth allows for the identification of enriched pathways associated with each target using conventional methods such as GSEA. Furthermore, having two treatment conditions allowed for the identification of targets exerting DE changes in one condition but not the other (Figure 2C-D). This highlights the importance of studying perturbations in relevant cellular contexts to uncover state-dependent regulatory effects, which are complex and poorly understood.

For early target discovery and the task of nominating promising targets for deeper investigation and investment, having ground truth for model evaluations is not as objective or clear as later parts of the drug discovery pipeline, such as predicting binding affinity between a small molecule and a target, or predicting cellular toxicity of a treatment. If a therapeutically efficacious target was known beforehand, then the efforts of target nomination are moot. However, as with most unmet disease areas, the efficacious targets to perturb are unknown. Hence, we need better predictions that are directly informed by the phenotype we are trying to achieve, which is the basis that motivated a latent similarity approach. More accurate target nomination, which is the beginning of a pipeline costing hundreds of millions of dollars for even one drug, can have substantial repercussions to costs and timeline of delivering efficacious medicines. Most early target nominees will fail *in vivo*, but if we can increase the success rate by even a fraction or gain greater biological understanding, then this means lower costs, faster delivery of therapies, and eventually more patient lives impacted.

A high enrichment score in the positive-control set indicated that the top ML-prioritized targets, which were prioritized solely by their transcriptomic signatures, were enriched with targets known to be relevant to the canonical TNF-α and IL-1β inflammation pathways and with anti-inflammatory potential. Having these positive-control targets is essential to evaluating the sensibility and potential for other neighboring targets in an ML-derived priority list. Their detection early in the priority rankings despite having no prior information about their relevance advocates for the feasibility of ML-guided target nomination. If positive-control targets were independently enriched in a ML-based ranking, as we observed in Figure 3C, then it begs the question: could other targets close in rank to the positive-control ones be a gap in our biological understanding and a potentially efficacious target with similar phenotypic effect?

While the DE (Inflammatory) ranking performed strongly in enrichment, the DE (Basal) ranking performed poorly (close to what was achieved by latent similarity via UMAP and Random representations) (Figure 3C). The poor DE (Basal) results demonstrate how using data generated under just basal conditions resulted in difficulty identifying gene targets related to specific disease states or conditions, hence advocating for the utility of having a complementary disease-relevant condition. While having multiple conditions for the same set of perturbations is more expensive, one key finding from this work is that knockdowns can have highly variable effects on the transcriptome depending on cellular state. Hence, the increase in cost is balanced with a closer alignment to disease-relevant biology. Based on the enrichment results, we advocate for target triaging in contexts that most closely resemble the disease state.

ChatGPT’s ranking of targets resulted in high AUC for the positive-control set (Figure 3C) despite having no access to our transcriptomic data and being prompted with just an English query summarizing the target nomination goal. High performance is perhaps not that surprising since ChatGPT was directly informed by human knowledge and literature trained on all the positive-control targets, which all have strong linkage to inhibition of TNF-α and IL-1β-driven inflammatory signaling. The latent similarity approaches based on scGPT and the raw transcript counts exceeded ChatGPT’s enrichment performance, despite never being informed about the positive-control set and its associations with the TNF-signaling cascades in the literature. Indeed, the latent similarity approach was based solely on an unbiased ranking of transcriptomic data, did not have access to the target labels, and simply transformed the raw transcriptome into representations without knowing which gene was perturbed. The latent similarity approach is uninformed by existing human knowledge and language, yet can create meaningful rankings that reflect known biology, which adds credibility to its use in novel target discovery. However, latent similarity based on STATE and SCimilarity had lackluster performance in this task (Figure 3C). Hence, it is important to note that the choice of representation has a large impact on performance, and not all target representations from the latent similarity approach led to high enrichment for the positive-control set. Surprisingly, the very simple approach of latent similarity via raw integer transcript counts as representations exceeded performance of both STATE and SCimilarity representations. This revealed the variability of different scFMs, and showed that scFMs can be exceeded by simpler methods depending on the task. The UMAP representation performed poorly, likely since it represented the entire transcriptome with just two dimensions, which is likely severely under-sized. However, larger representation sizes did not necessarily mean higher performance, with lower-sized embeddings like scGPT (size = 512) outcompeting representations with much larger sizes (STATE size = 2,058 and Raw size = 38,606).

The latent similarity approach also resulted in a target ranking that was significantly enriched for relevant pathways chosen *a priori*. scGPT in particular resulted in 100% recall of the relevant pathways for its top 30 to top 100 targets (Figure 4A). scFMs such as scGPT do not have any concept of pathways, have no way of being informed about the relevance of the *a priori* chosen ones, and ranked targets solely on transcriptomic similarity to control data unperturbed by TNF-α and IL-1β (as opposed to ChatGPT which is directly informed by the literature about different pathways). Hence, the fact that the latent similarity approach could construct a target ranking that can recall relevant pathways of interest is evidence of scFMs’ propensity to create biologically meaningful target rankings based solely on their internal learned representations applied to transcriptomic data. We would expect the top targets from a meaningful target ranking to be directly related to TNF-α and IL-1β signaling pathways, and this was confirmed by the Enrichr results (Figure 4A). When queried against a larger library outside of the *a priori* chosen pathways, we also observed sensible pathways that are directly related to TNF-α and IL-1β signaling (Figure 4B). scFMs have no direct concept of pathways. Hence, the fact that relevant pathways can be independently surfaced via an unbiased ranking of the transcriptomic latent space adds some confidence to the biological relevance of triaged targets. Since the top targets from the DE-approach rankings did not yield significantly enriched pathways of apparent relevance (Figure 4B), this could advocate for greater sensibility of target rankings produced by latent similarity over rankings from a DE approach.

For both target enrichment (Figure 3) and pathway enrichment (Figure 4), ChatGPT and the front-running representations in the latent similarity approach both had higher performance than the DE-approach rankings, advocating for the potential use of ML for target triaging. Since they performed better than DE-approach rankings on relevant target and pathway enrichment, perhaps the targets highly prioritized in these rankings but not in the DE-approach rankings are candidates worth further exploration for the sake of novel discovery.

Several caveats limit and focus the scope of this study. Our validation consisted primarily of assessing enrichment of known targets and pathways, and did not include any *in vitro* or *in vivo* follow-up of highly prioritized targets, which would be the only way to definitively validate a target ranking. Also, as with all early target discovery programs, target druggability and potential pleiotropic effects can completely inhibit progression in the drug development pipeline, regardless of any promising phenotype reversion induced by *in vitro* target inhibition. Prioritizing targets solely by phenotype in latent space cannot possibly account for these issues and may suggest candidates with no possible therapeutic promise. Furthermore, our experiment’s model system of TNF-α and IL-1β induced inflammation might not accurately capture the actual *in vivo* phenotype of diseased states, and hence any targets derived from this system may not be relevant to treating disease in humans. *In vivo* validation in a system more closely approximating human disease is necessary for any clinical viability, which our study does not address.

Lastly, although the positive-control set has strong literature backing, it is innately biased and based on the expertise of biologists working in the space. If a different set were chosen, it could shift our choice of algorithm for follow up analyses. Due to the lack of objective ground truth in the early target discovery phase, this is an accepted limitation that focuses the scope in assessing the quality of target triaging. Additionally, having only six positive-control targets is too small of a sample to make definitive claims for triaging. However, the fact that these positive-control targets were enriched in ML-based rankings derived with no access to the literature demonstrates a convergence between human-derived insights based on current knowledge and data-driven ML insights based solely on data.

Despite clear limitations of a latent similarity approach for early target discovery, there are advantages over a DE approach. The DE approach relies heavily on relevant pathway selection, which itself is limited and does not contain pathways that are specific enough to mirror the precise phenotype of interest given our cellular context. Annotated gene sets are generalized pathways that cannot possibly describe the reality of our specific biological context. They lack the granularity of differential gene function in specific cell-types, which can be highly variable. The DE approach also relies heavily on good pathway annotations, which can be sparse, heavily biased towards genes that are well studied, and is constantly evolving. Contrast this with a data-first approach like latent similarity that does not depend on pathway annotations or humans selecting what they believe to be relevant pathways to make target rankings. Furthermore, The TNF-α and IL-1β pro-inflammatory signaling that was explored as the stimulation of interest in this dataset has been well-studied and has a wealth of pathway annotations. These annotations and pathway gene sets can aid in the DE approach to identify genes reversing the inflammatory phenotype. However, for other phenotypic stimulations such as flow response or oxidized low-density lipoprotein exposure, which are both processes important for plaque development, the pathway annotations are not well developed or curated. In this situation, the DE approach would be much more difficult to use for identification of genes reversing the phenotype. Yet the latent similarity approach could offer unique opportunities in sparsely studied and annotated situations like these. The latent similarity approach benefits by not being dependent on the presence or quality of pathway annotations. Therefore, it can be used to investigate genes driving expression changes related to any stimulation. Pathway annotations themselves are biased to current biological understanding and the interpretation of the curators, while the latent similarity approach does not rely on such manual interpretations. It requires less user input and is less biased, which can be advantageous for novel discovery.

The latent similarity approach also has advantages over English-based LLMs like ChatGPT. LLM-based tools are incredibly useful for summarizing and linking different facets of human knowledge, but suffer the same bias of the DE approach in that they depend heavily on current human knowledge and are biased towards aspects of biology that have a large publication record, which was used for training ChatGPT. Such a bias seems at odds with novel discovery. Also, these English-based LLMs cannot be easily translated to operate on the transcriptomic counts of RNA-seq data, making it difficult to apply the concepts learned during pre-training to our specific data modalities. Latent similarity based on scFMs are designed to operate directly on the “language” of the transcriptome and hence are better equipped to operate on the RNA-seq datasets so common in early discovery programs. Although scFMs have their own issues and biases, the latent similarity approach is less biased by human intuition and literature and hence may be favorable for discovery beyond the current state of human knowledge.

The general approach of deriving meaningful representations for entities like perturbations or basal cell states and then querying those representations in a high dimensional latent space is a model-agnostic framework. This is incredibly important as new foundation models are continually developed and improved at a dizzying pace in the field. Latent similarity analyses can be re-run with improved representations, with no new training or fine-tuning required—just forward passes with pre-existing models which have quick inference times. Therefore, the approach does not require computationally expensive resources like those for training new models end-to-end. Aligned with a core principle in ML of data-driven insights, this latent similarity approach of ranking targets is based solely on modeling phenotypes of interest (e.g. the transcriptomic signatures of perturbations) with little human bias or prior biological knowledge required. Although expert human knowledge is invaluable, this primarily data-driven approach can add a unique arm of evidence to target nomination campaigns that can supplement more traditional analyses centered on human knowledge.

In summary, these findings yield several important takeaways: (1) Perturb-seq datasets with both a stimulated condition and a basal condition add great value in identifying targets driving changes in gene expression related to disease-relevant stimulation. (2) The latent similarity approach can identify biologically-relevant genes related to the condition of interest as well as or better than the DE approach and a powerful English-based LLM like ChatGPT, all without the burden and bias of relying on human-derived pathway annotations or literature. As a future direction, we hope to perform validation on a subset of the top ML-ranked targets. Confirmation via target and pathway enrichment is an encouraging but preliminary result, but the only way to verify them is via focused study and inquiry on high priority targets via *in vivo* studies. We hope that this study’s proof-of-concept can serve as the basis for other phenotypic reversion and target nomination campaigns for other disease-associated phenotypes of interest. We also hope that the release of the Perturb-seq dataset will facilitate new biological inquiry and discovery both through direct inquiry and improved ML capabilities.

## Methods

### Creation of Perturb-seq dataset

#### Cell Culture

TeloHAEC cells were obtained from ATCC and cultured in Endothelial Cell Growth Medium-2 (EGM-2, Lonza) and EGM-2 BulletKit (CC-3162, Lonza). To induce the cells into an inflammatory state, the cells were treated with 1ng/mL TNF-α and 1ng/mL IL-1β for 24 hours. A CRISPRi dCas9-KRAB vector with an mCherry selection marker and EFS promoter (Cellecta) was used to generate lentivirus. 5×10^6^ cells were spinfected with the CRISPRi virus at 2,000 rpm for 2 hours at 30C with 10μg/mL polybrene. These cells were then sorted twice on a Sony Cell Sorter to establish a cell population of >95% mCherry positive cells.

#### Generation of pooled dual-sgRNA lentiviral library

Each gene was targeted by three constructs within the CRISPRi library, and each construct contained paired sgRNA sequences targeting the same gene. Paired sgRNAs were selected from the Replogle^39^ and Dolcetto^40^ libraries, with priority given to the Replogle sgRNAs. While the Replogle designs were already paired, the Dolcetto library consists of single sgRNAs. The sgRNAs in the Dolcetto library were paired as 1+4, 2+5, and 3+6, with 1 being the top ranked sgRNA, with the exception that sgRNAs also present in the Replogle library were skipped. The paired sgRNAs were cloned into the pWRS1001_U6_H1_PuroR dual sgRNA lentiviral expression vector^41^ with a GFP selection marker in two sub-pools. In addition, safe-harbor guides^42^ and non-targeting guides^40^, which served as negative controls, were selected and paired at random within each negative control class. 36 safe-harbor pairs and 36 non-targeting pairs were included in each sub-pool.

#### TeloHAEC CRISPRi perturb-seq screen

Equal amounts of DNA from each sub-pool were combined and used to generate pooled lentivirus expressing dual-sgRNAs for CRISPRi. 12 x10^6^ TeloHAEC CRISPRi cells were spinfected with the dual-sgRNA virus at 2000rpm for 2 hours at 30°C with 10ug/mL polybrene at a 0.3 MOI. Three days after infection, cells were sorted for expression of both GFP (dual-sgRNA vector) and mCherry (CRISPRi dCas9-KRAB vector) and cultured for an additional 2 days. 6×10^6^ sorted cells were then seeded for each of the two treatment conditions. After allowing one day for the cells to adhere, cells were either treated with 1ng/mL TNF-α and 1ng/mL IL-1β for 24 hours (inflammatory condition) or left untreated with DMSO (basal condition). After treatment, cells were harvested from each condition and subjected to single-cell RNA sequencing. Treated and untreated cells were processed in parallel, with eight samples of each condition processed per batch.

Single-cell transcriptomic profiling was performed using the Chromium Next GEM Single Cell 5′ HT Kits V2 according to the manufacturer’s protocol. Single-cell suspensions were prepared from freshly isolated 2.4 x 10^6^ cells per condition (treated vs untreated) and filtered through a 30-µm cell strainer to remove debris and aggregates. Cell viability was assessed using trypan blue exclusion, and samples with >85% viable cells were further processed.

Approximately 0.8 x 10^6^ of cells per sample were loaded onto the Chromium Next GEM Chip N to achieve a targeted recovery of 5,000 cells/µl per channel. Individual cells were encapsulated with barcoded gel beads in oil droplets to generate Gel Bead-in-Emulsions (GEMs), enabling cell-specific barcoding and unique molecular identifier (UMI) tagging during reverse transcription. After GEM generation, reverse transcription, cDNA amplification, and library construction were performed according to the 10x Genomics 5′ gene expression workflow.

Final libraries were quantified using a Qubit fluorometer and assessed for fragment size distribution using a Bioanalyzer. Pooled scRNAseq libraries were sequenced on a Novaseq S4 sequencer (Illumina). Raw sequencing files were processed with CellRanger v8.0.1 (10X Genomics) to generate a cell x transcript expression read count matrix.

#### Demultiplexing, alignment, and quantification of guide and gene expression reads

CellRanger count (v8.0.1) was run for demultiplexing, alignment and quantification of gene expression and guide reads by specifying the Single Cell 5′ R2-only chemistry (SC5P-R2). Cellranger count was performed for each of the numbered wells and combined the counts across flow cells for each well. Since there were two conditions present, unstimulated and TNF-α and IL-1β stimulated conditions, we performed CellRanger Aggr (v8.0.1) to aggregate the counts across all numbered wells for each condition. Default parameters were used for CellRanger Aggr, except no normalization for sequencing depth was performed. The filtered count matrix by CellRanger Aggr was utilized for downstream methods.

#### Assignment of guide RNAs to cells

Guide identification was initially performed using CellRanger (v8.0.1), which fit a Gaussian Mixture Model to the log-transformed guide RNAs / cell distribution for each guide RNA. Of the 1,750 genes targeted with guide RNAs, 1,745 genes had guides assigned. All of the guide sequences can be found in Supplemental Table 1.

Upon inspection of the number of guide UMIs mapping to each cell, we found a bimodal distribution, with background levels containing less than 13 guide UMIs per cell. We filtered the cells with less than 13 guide UMIs to remove the background consisting of non-guide-containing droplets. Next, we selected for only those cells containing guides from a single gene perturbation. To ensure robust selection of single gene-targeted cells, a cell was defined as having a single perturbation if the gene receiving the largest number of guide UMIs was greater than or equal to four times the number of UMIs for the gene receiving the next largest number of guide UMIs, adapted from the method used by Schnitzler et. al^42^. All cells not identified as containing a single gene target were filtered from the dataset leaving 887,539 cells.

#### scRNA-seq gene expression quality filtering

Quality metrics were explored for the scRNA-seq gene expression data, including number of UMIs (nUMIs), number of genes (nGenes), and mitochondrial content. The following thresholds were applied for quality filtering: 1200 nGenes, 2500 nUMIs, 0.8 log_10_ genes per UMI (log_10_(nGenes)/log_10_(nUMIs)), and less than 20% mitochondrial reads. Filtering was performed using ScanPy (v1.11.1), and after filtering, there were 864,115 remaining cells.

### Dataset analysis

We calculated KD efficiency according to the method presented in Replogle et al. for a cell with target = G and treatment condition C as follows:

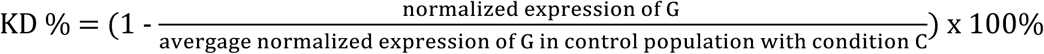

For normalization, we had each cell’s total transcript count sum to the median UMI. We aggregated cells by their target and averaged over targets to get the final reported KD efficiency. When we performed a UMI filter on the genes and removed any targets that had an average UMI less than 1.0 across all cells, we had a KD efficiency mean of 70% and median of 78%.

Differential expression of genes was calculated as follows. For every perturbed gene, the Wilcoxon rank-sum test was employed to determine the genes differentially expressed between the cells receiving a particular perturbation and the control cells (including both safe-targeting and non-targeting). To determine significant differentially expressed genes, we used a Benjamini-Hochberg p-adjusted threshold of 0.05.

For Supplemental Figure 5, we assessed same-target replicate similarity by comparing cell clustering by the true gene-target label vs cell clustering by a randomized label. This was performed for each of the two treatment conditions independently using all the gene target labels (excluding the safe-targeting and no-targeting controls). We used the calinski_harabasz_score function from scikit learn.

### Deriving the different rankings for each approach

#### See the GitHub repository for all source code pertaining to ranking construction

The rankings for the DE (Basal) approach were derived as follows. For each target, we performed Gene set enrichment analysis (GSEA) using the fgsea package (v1.28.0) using R v4.3.2. Genes were ranked using signed log-transformed p-values ( sign(log(fold-change))*-log_10_(p-value)) from the DE analysis results, and tested against the following six MSigDB C2 canonical pathways: BioCarta: IL1R Pathway, BioCarta: TNFR1 Pathway, KEGG: Cytokine-Cytokine Receptor Interaction, Reactome: Interleukin 1 Family Signaling, Reactome: Interleukin 1 Signaling, Reactome: TNF Signaling. Pathways and their corresponding gene sets were pulled from v7.0 of the MSigDB C2 canonical pathways released on Aug 2019. We used the standard adjusted p-value < 0.05 to determine significance. fgsea was run with default parameters (min set size = 1 and max set size = number of tested genes - 1). Next, we ranked the targets by the number of significant down-regulated pathways, with the most number of significant down-regulated pathways as the highest priority. If two targets had an equal number of significant down-regulated pathways, we prioritized the one with the lowest adjusted p-value of the most significant down-regulated pathway.

For the DE (Inflammatory) approach, we followed a similar process, but with an additional step that deprioritized targets with significant down-regulated pathways in both the basal and inflammatory conditions, since these pathways were less specific to the inflammatory condition. These targets with down-regulated pathways in both conditions were included after the inflammatory-specific condition targets, and ranked by p-value and direction of regulation similar to the targets not significantly enriched for any of the inflammatory pathways.

For the latent similarity approach, we performed neither pre-processing on the data nor filtering, and used each individual method’s default pre-processing pipelines prior to running a forward pass on each cell. We used the following pre-trained models to create the high-dimensional representations for each cell: SCimilarity (v0.1.0), scGPT (v0.2.1), and STATE (v0.9.3). For scGPT, we used the “scGPT_CP” pre-trained model. All models were run zero-shot with no fine-tuning. The “Raw” method simply used the raw integer counts and the entire transcriptomic space as the feature embedding. The UMAP representation was calculated by performing a two-dimensional UMAP reduction over the raw expression matrix with all transcriptomic features included.

After embedding each cell into a high-dimensional representation, we constructed perturbation-level representations (separately for the treated and untreated conditions). For each target gene G and treatment condition C, the aggregated representation was calculated by averaging over every cell’s representation with target gene = G and treatment = C. Each method’s latent representation was relatively stable with differing cell counts (Supplemental Figure 7).

Once representations for each perturbation target were calculated, we computed cosine similarities between the untreated safe-targeting representation and each treated perturbation target. We chose the safe-targeting control (instead of the no-targeting control) as our desired phenotype because it most resembled the treated cells perturbed by CRISPRi. This served to control for spurious background factors related to the CRISPR dCas9 mechanism (which no-targeting does not engage) and to focus instead on differences in gene expression inhibition between the different perturbations. Perturbations were ordinally ranked from highest priority to lowest based on highest cosine similarity (to the control) to lowest.

For the ranking derived by ChatGPT, we used the following prompt: “You are an expert biologist with deep understanding of signaling cascades. I ran an experiment where I introduced TNF-α and IL-1β cytokines to teloHAEC cells. I also introduced each of the following gene targets to knockdown via CRISPRi. Which knockdowns are the most likely to induce a transcriptomic state that most resembles the normal cell state untreated with the cytokines? Rank my entire list by most likely to induce a shift to the normal cell state to least likely given my experimental setup, and return it to me as a downloadable CSV.”

### Evaluating rankings via target enrichment

For the positive-control gene set, we gave an internal set of experts the biological context of the Perturb-seq experiment, a list of all perturbations, and asked them to nominate targets that would induce an anti-inflammatory phenotype upon CRISPRi treatment. This was performed independently of all analyses, and the experts did not have any prior knowledge about any rankings. The following set of genes was given: TNFRSF1A (a major TNF-α receptor), TRADD (a major TNF-α signaling adaptor), and the following major transcriptional factors that mediate inflammatory gene expression: JUNB, JUND, NFKB1, NFKB2.

We computed enrichment scores by calculating the AUC based on the sorted ranking indices vs the indices at which a reference gene was placed in the ranked list. An AUC of 100% indicates perfect enrichment in which all the positive controls are placed at the top of the priority queue. For all the gene sets based on pathway annotations (Supplemental Figure 6), we filtered the sets to only include targets that were present in our study prior to computing enrichment.

### Evaluating rankings via pathway enrichment

For the library-based pathways in Figure 4A, we chose the same six relevant pathways we used to create the DE-approach rankings. Pathways and their corresponding gene sets were pulled from MSigDB C2 canonical pathways (v7.0) released on Aug 2019. For the broader pathway enrichment analysis in Figure 4B, we computed significance for each pathway in the following five libraries from the GSEAPY library via Enrichr: KEGG_2021_Human, BioCarta_2016,Reactome_Pathways_2024, MSigDB_Hallmark_2020, and GO_Biological_Process_2025. For the background set of genes, we used all the target genes in our study, and a false-discovery-rate-adjusted p-value of 0.05 as the significance threshold.

## Supplemental Figures

**Supplemental Figure 1:**
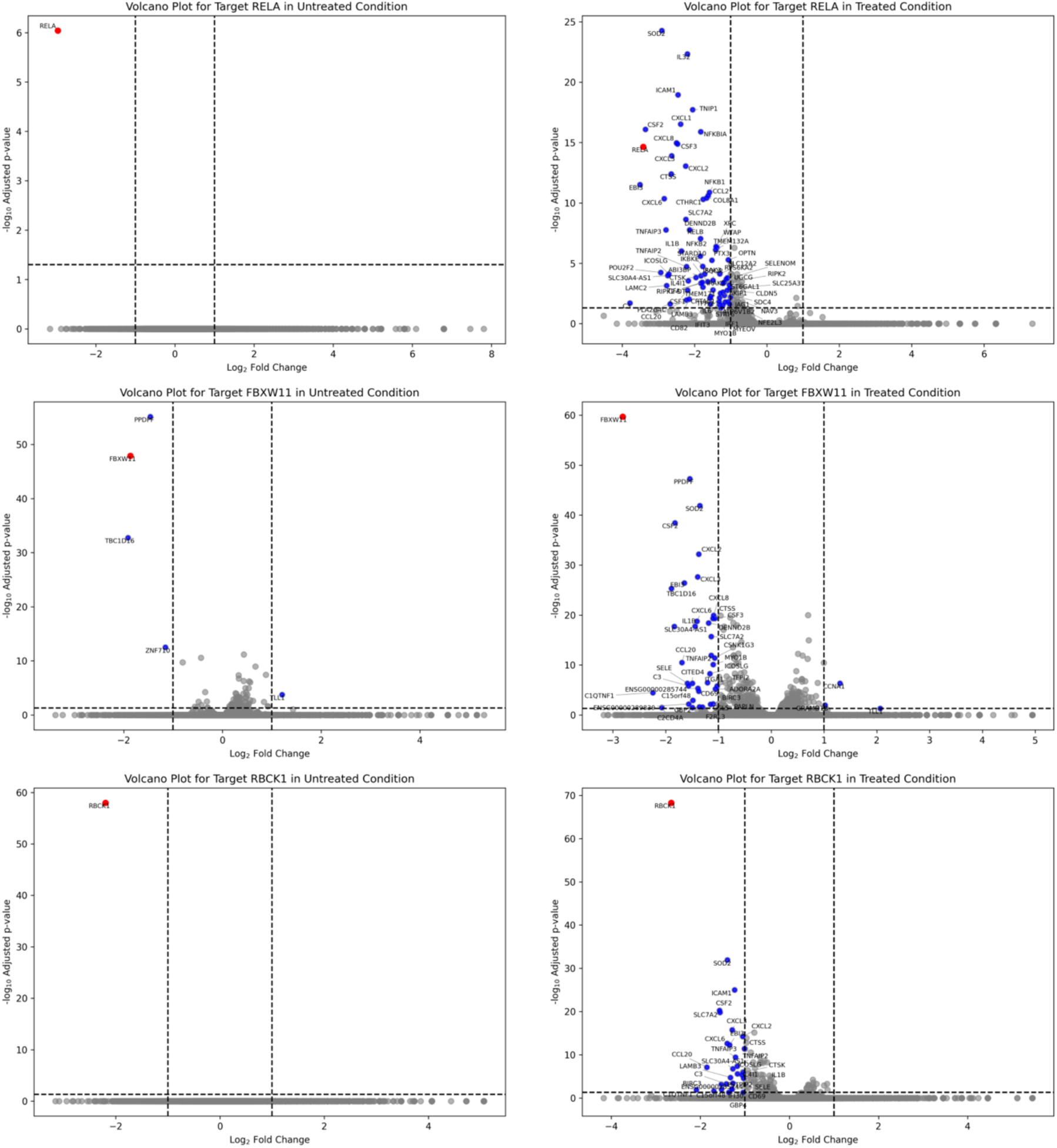
Volcano plots for example targets with differential effect in treated vs untreated conditions.

**Supplemental Figure 2:**
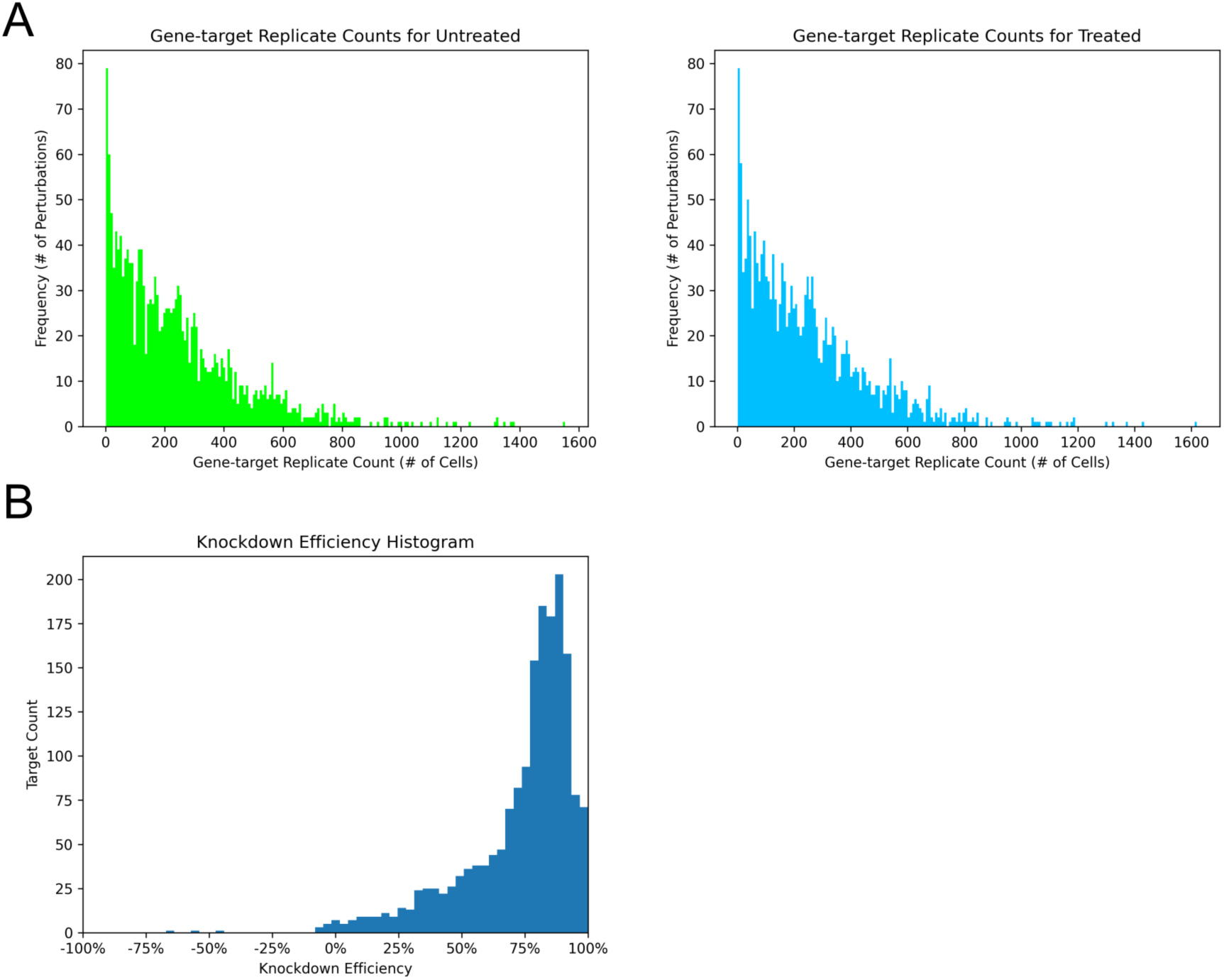
distribution of gene target replicate counts and knockdown efficiency. (a) Histogram of gene target replicate counts for each condition (excluding the safe-targeting and no-targeting controls). (b) Histogram of knockdown efficiencies for each target. X-axis is limited to -100% to 100%.

**Supplemental Figure 3:**
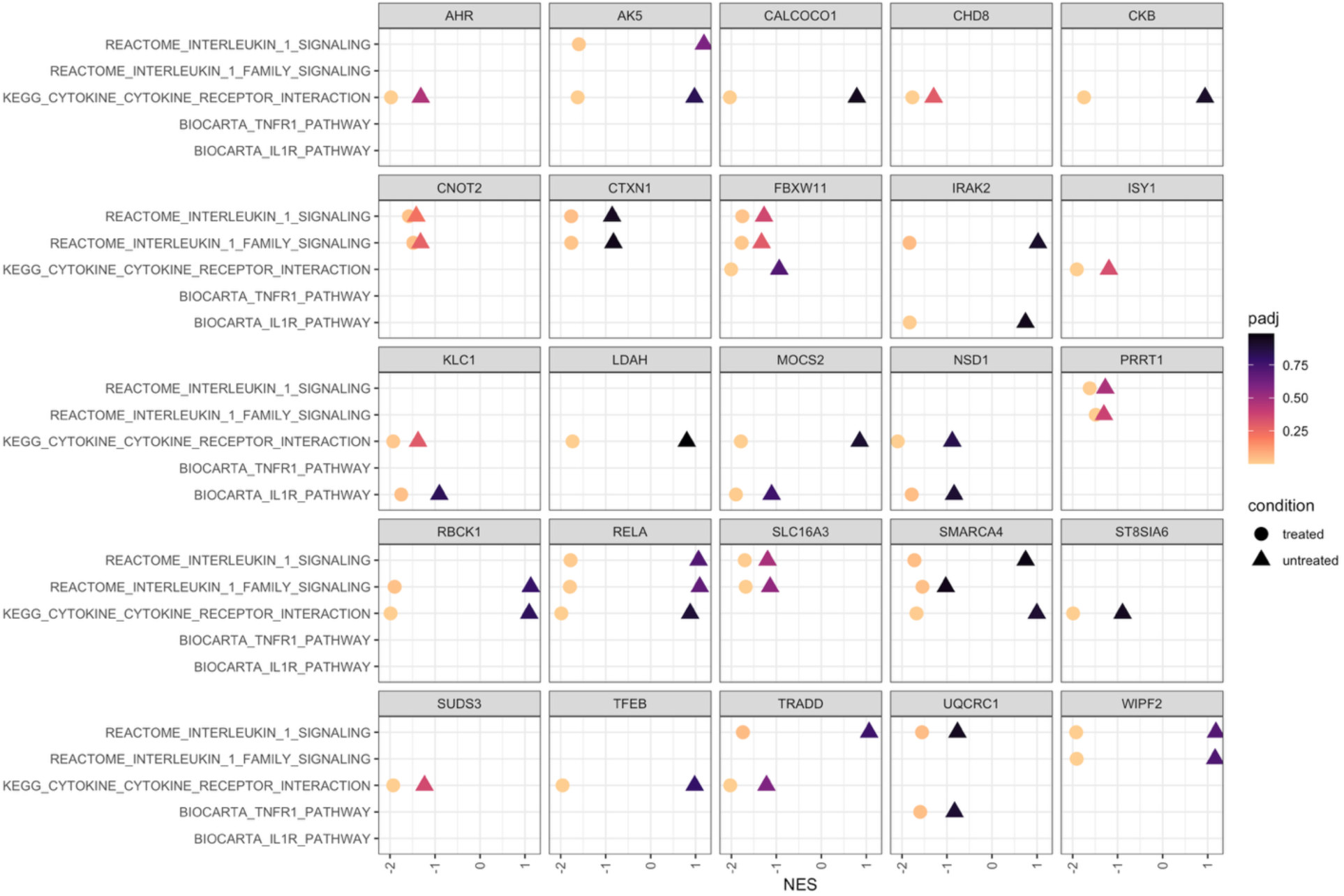
Top 25 targets for the DE (Inflammatory) ranking. X-axis normalized enrichment score returned from GSEA. Y-axis: relevant pathways. The “Reactome: TNF Signaling” pathway was not significantly enriched for any of the targets and hence is not shown on the Y-axis.

**Supplemental Figure 4:**
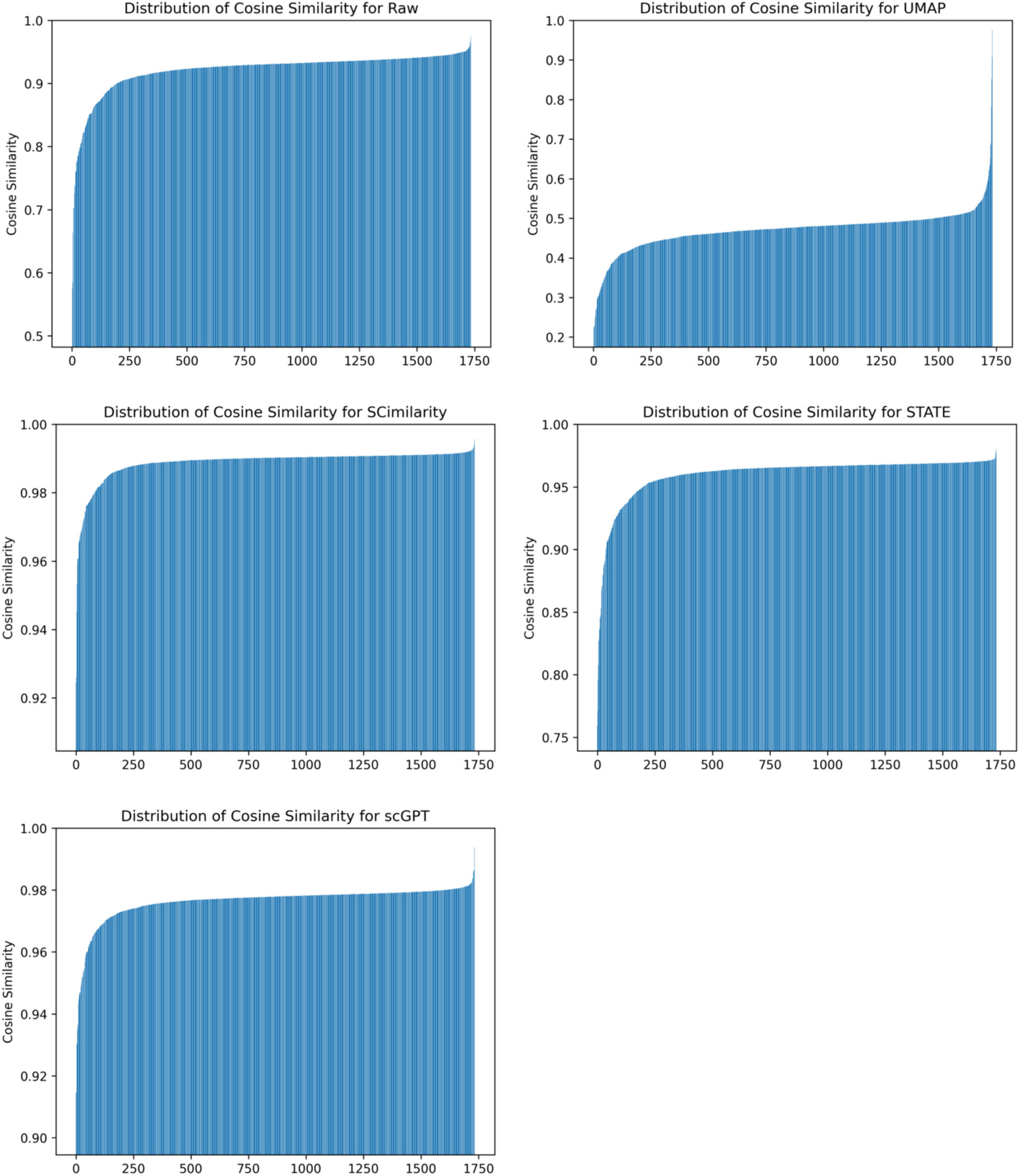
Distribution of sorted cosine similarities for the different methods of the latent similarity approach.

**Supplemental Figure 5:**
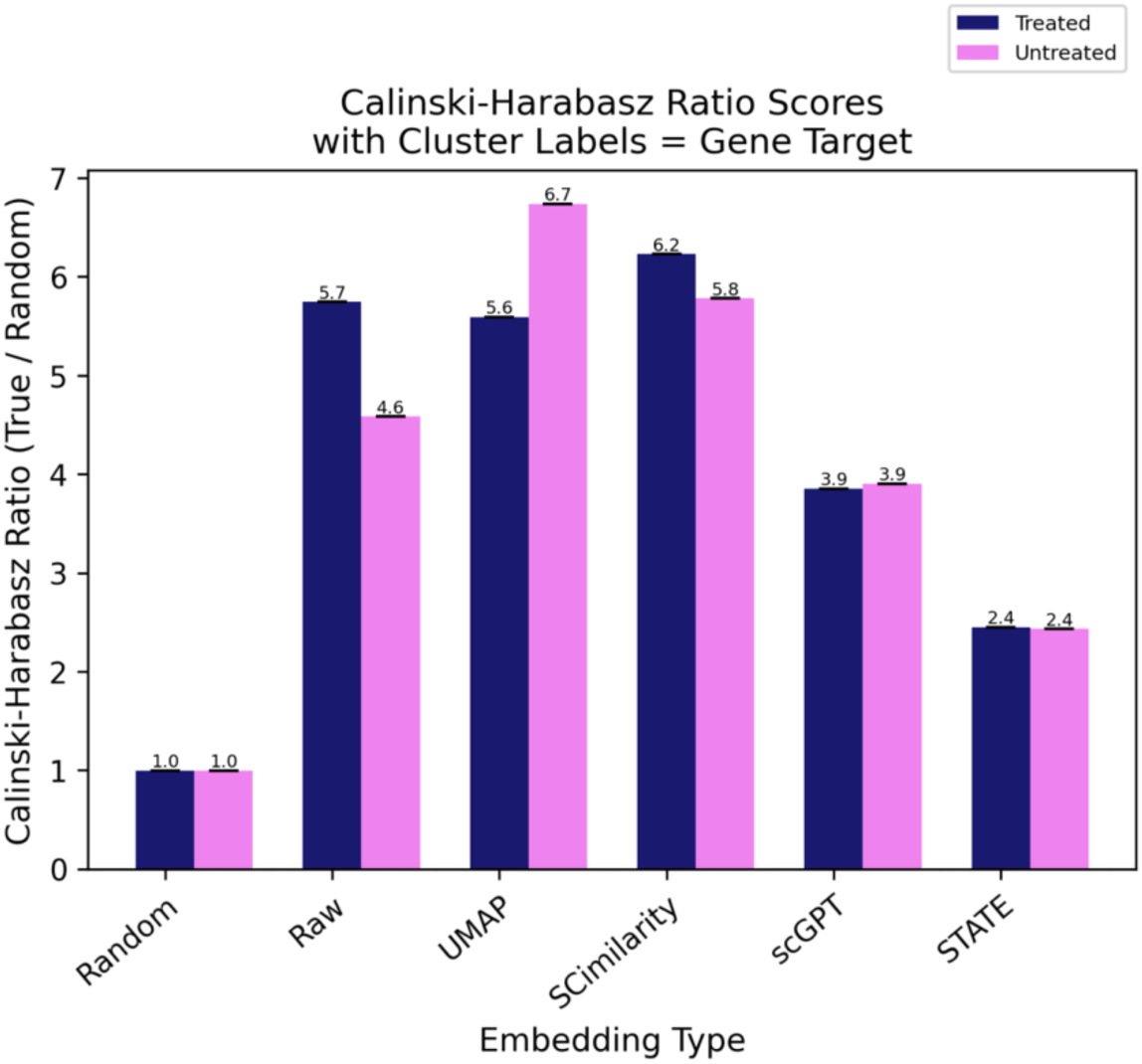
Clustering embeddings of the latent similarity approach with biologically known labels. Calinski-Harabasz scores when assigning cluster labels as the true gene target labels divided by the scores when assigning randomly shuffled target labels. Control cells were removed from this analysis to focus on clustering of genetic targets.

**Supplemental Figure 6:**
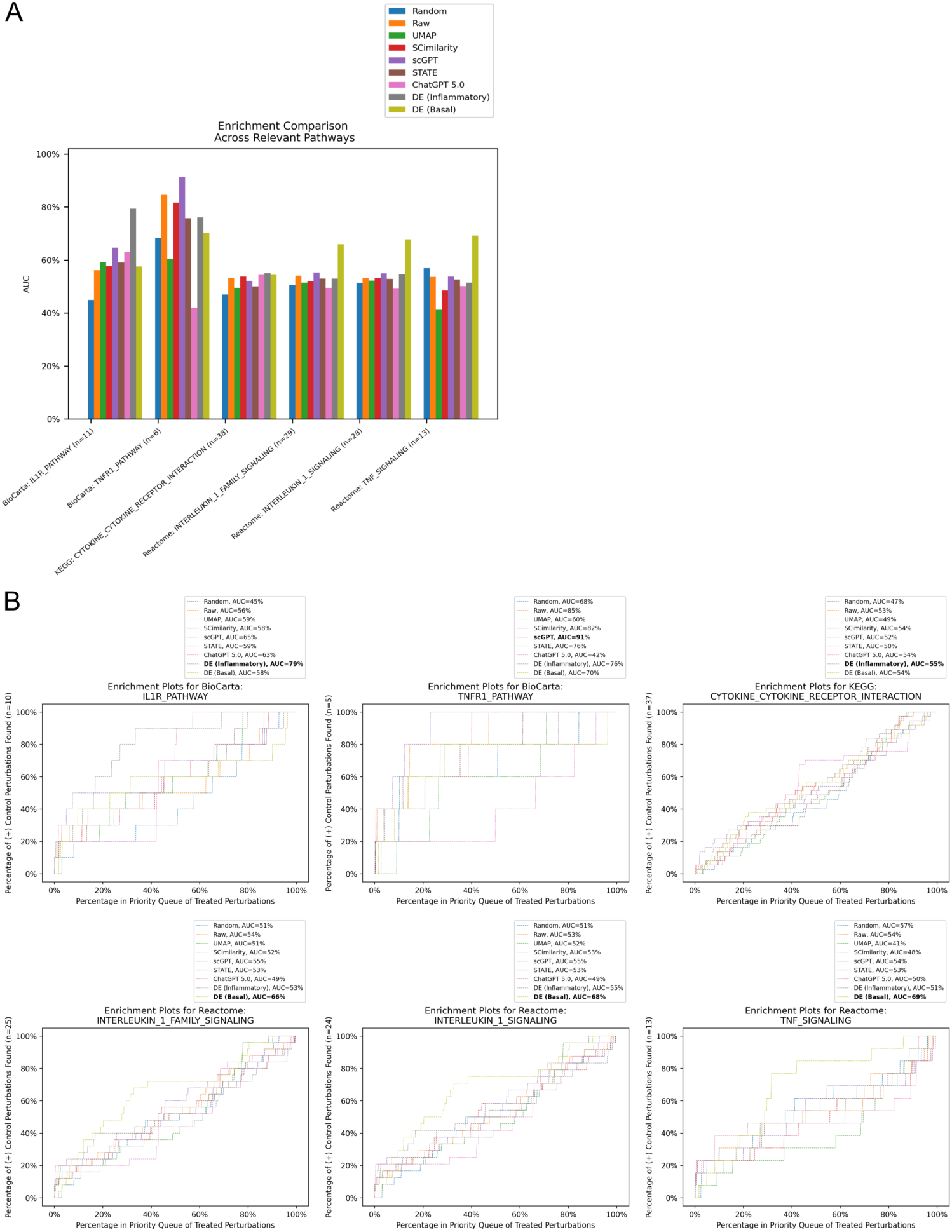
Enrichment AUC for relevant annotated pathways. (a) X-axis indicates the reference pathway and gene set. N indicates the number of genes part of the enrichment set. For all pathways, we subselected to just the gene targets in our dataset. Y-axis: AUC of enrichment. (b) Enrichment plots for each pathway. The bolded method in the legend is the one with the highest AUC.

**Supplemental Figure 7:**
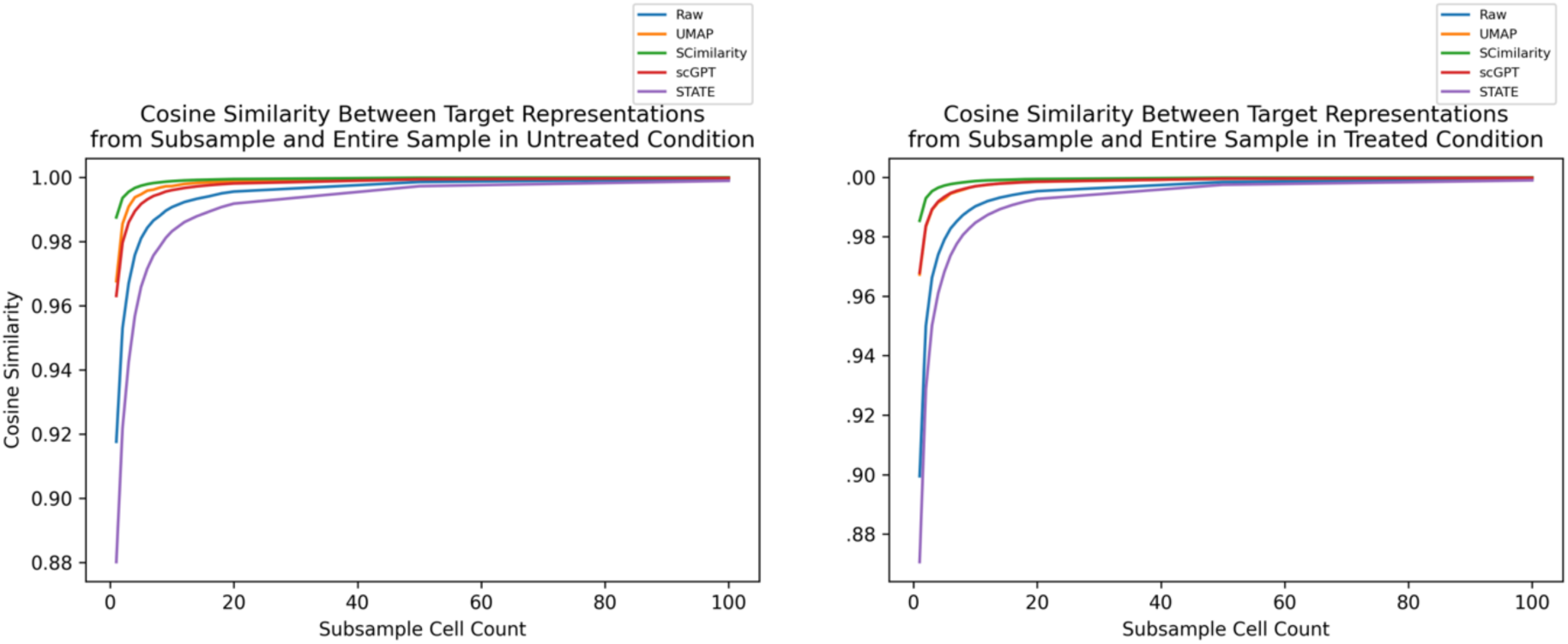
Cosine similarity between full sample latent representation and subsampled latent representations. Each point in this plot is the average cosine similarities between target T_A_ and target T_S_ for all T in the target set such that T_A_ is the target representation derived from all cells of target T, and T_S_ is the target representation derived from a subset of cells of target T sampled at random without replacement. The sample size is determined by the x-axis.

## Data Availability

All datasets are public and can be found at DOI: 10.5281/zenodo.18792213. All source code is available at: https://github.com/pfizer-opensource/phenotype_reversion.

## Acknowledgements

We thank Margery Pelletier and Vaughn Youngblood for providing the resources necessary for flow cytometry.

## References

1. Lowe, R., Shirley, N., Bleackley, M., Dolan, S. & Shafee, T. Transcriptomics technologies. PLoS Comput. Biol. 13, e1005457 (2017).

2. Tzec-Interián, J. A., González-Padilla, D. & Góngora-Castillo, E. B. Bioinformatics perspectives on transcriptomics: A comprehensive review of bulk and single - cell RNA sequencing analyses. Quant. Biol. 13, (2025).

3. Molla Desta, G. & Birhanu, A. G. Advancements in single-cell RNA sequencing and spatial transcriptomics: transforming biomedical research. Acta Biochim. Pol. 72, 13922 (2025).

4. Tang, F. et al. mRNA-Seq whole-transcriptome analysis of a single cell. Nat. Methods 6, 377–382 (2009).

5. Tabula Sapiens Consortium* et al. The Tabula Sapiens: A multiple-organ, single-cell transcriptomic atlas of humans. Science 376, eabl4896 (2022).

6. Jinek, M. et al. A programmable dual-RNA-guided DNA endonuclease in adaptive bacterial immunity. Science 337, 816–821 (2012).

7. Dixit, A. et al. Perturb-seq: Dissecting molecular circuits with scalable single-cell RNA profiling of pooled genetic screens. Cell 167, 1853–1866.e17 (2016).

8. Chen, B. et al. Reversal of cancer gene expression correlates with drug efficacy and reveals therapeutic targets. Nat. Commun. 8, 16022 (2017).

9. Wong, D. R. et al. Deep representation learning determines drug mechanism of action from cell painting images. Digit. Discov. 2, 1354–1367 (2023).

10. Bray, M.-A. et al. Cell Painting, a high-content image-based assay for morphological profiling using multiplexed fluorescent dyes. Nat. Protoc. 11, 1757–1774 (2016).

11. Kraus, O., et al. Masked autoencoders are scalable learners of cellular morphology. arXiv [cs.CV] (2023).

12. Wong, D. R. et al. Trans-channel fluorescence learning improves high-content screening for Alzheimer’s disease therapeutics. *Nat*. Mach. Intell. 4, 583–595 (2022).

13. Page, M. J., Amess, B., Rohlff, C., Stubberfield, C. & Parekh, R. Proteomics: a major new technology for the drug discovery process. Drug Discov. Today 4, 55–62 (1999).

14. Scaffidi, P. & Misteli, T. Reversal of the cellular phenotype in the premature aging disease Hutchinson-Gilford progeria syndrome. Nat. Med. 11, 440–445 (2005).

15. Wagner, A. et al. Drugs that reverse disease transcriptomic signatures are more effective in a mouse model of dyslipidemia. Mol. Syst. Biol. 11, 791 (2015).

16. Carvalho, R. F. et al. Drug repositioning based on the reversal of gene expression signatures identifies TOP2A as a therapeutic target for rectal cancer. Cancers (Basel) 13, 5492 (2021).

17. Plesa, A. M., Shadpour, M., Boyden, E. & Church, G. M. Transcriptomic reprogramming for neuronal age reversal. Hum. Genet. 142, 1293–1302 (2023).

18. Bommasani, R., et al. On the opportunities and risks of foundation models. arXiv [cs.LG] (2021).

19. Cui, H. et al. Towards multimodal foundation models in molecular cell biology. Nature 640, 623–633 (2025).

20. Cui, H. et al. scGPT: toward building a foundation model for single-cell multi-omics using generative AI. Nat. Methods 21, 1470–1480 (2024).

21. Heimberg, G. et al. A cell atlas foundation model for scalable search of similar human cells. Nature 638, 1085–1094 (2025).

22. Theodoris, C. V. et al. Transfer learning enables predictions in network biology. Nature 618, 616–624 (2023).

23. Wong, D. R., Hill, A. S. & Moccia, R. Simple controls exceed best deep learning algorithms and reveal foundation model effectiveness for predicting genetic perturbations. Bioinformatics 41, (2025).

24. Boiarsky, R., Singh, N., Buendia, A., Getz, G. & Sontag, D. A deep dive into single-cell RNA sequencing foundation models. bioRxiv (2023) doi:10.1101/2023.10.19.563100.

25. Lopez, R., Regier, J., Cole, M. B., Jordan, M. I. & Yosef, N. Deep generative modeling for single-cell transcriptomics. Nat. Methods 15, 1053–1058 (2018).

26. Wu, J. et al. Biology-driven insights into the power of single-cell foundation models. Genome Biol. 26, 334 (2025).

27. Adduri, A. K., et al. Predicting cellular responses to perturbation across diverse contexts with State. bioRxiv (2025) doi:10.1101/2025.06.26.661135.

28. Hao, M. et al. Large-scale foundation model on single-cell transcriptomics. Nat. Methods 21, 1481–1491 (2024).

29. Kalfon, J., Samaran, J., Peyré, G. & Cantini, L. scPRINT: pre-training on 50 million cells allows robust gene network predictions. Nat. Commun. 16, 3607 (2025).

30. Barbadilla-Martínez, L., Klaassen, N., van Steensel, B. & de Ridder, J. Predicting gene expression from DNA sequence using deep learning models. Nat. Rev. Genet. 26, 666–680 (2025).

31. Libby, P. Inflammation in atherosclerosis. Nature 420, 868–874 (2002).

32. Pepin, M. E. & Gupta, R. M. The role of endothelial cells in atherosclerosis: Insights from genetic association studies. Am. J. Pathol. 194, 499–509 (2024).

33. Jiang, H. et al. Mechanisms of oxidized LDL-mediated endothelial dysfunction and its consequences for the development of atherosclerosis. Front. Cardiovasc. Med. 9, 925923 (2022).

34. Liu, T., Zhang, L., Joo, D. & Sun, S.-C. NF-κB signaling in inflammation. Signal Transduct. Target. Ther. 2, 1–9 (2017).

35. Mai, W. & Liao, Y. Targeting IL-1β in the treatment of atherosclerosis. Front. Immunol. 11, 589654 (2020).

36. Xie, Z. et al. Gene set knowledge discovery with Enrichr. Curr. Protoc. 1, e90 (2021).

37. Fang, Z., Liu, X. & Peltz, G. GSEApy: a comprehensive package for performing gene set enrichment analysis in Python. Bioinformatics 39, (2023).

38. Roohani, Y. H. et al. Virtual Cell Challenge: Toward a Turing test for the virtual cell. Cell 188, 3370–3374 (2025).

39. Replogle, J. M. et al. Mapping information-rich genotype-phenotype landscapes with genome-scale Perturb-seq. Cell 185, 2559–2575.e28 (2022).

40. Sanson, K. R. et al. Optimized libraries for CRISPR-Cas9 genetic screens with multiple modalities. Nat. Commun. 9, 5416 (2018).

41. Li, R. et al. Comparative optimization of combinatorial CRISPR screens. Nat. Commun. 13, 2469 (2022).

42. Schnitzler, G. R. et al. Convergence of coronary artery disease genes onto endothelial cell programs. Nature 626, 799–807 (2024).

